# Single-cell RNA-seq analysis of human coronary arteries using an enhanced workflow reveals SMC transitions and candidate drug targets

**DOI:** 10.1101/2020.10.27.357715

**Authors:** Wei Feng Ma, Chani J. Hodonsky, Adam W. Turner, Doris Wong, Yipei Song, Nelson B. Barrientos, Jose Verdezoto Mosquera, Clint L. Miller

## Abstract

**Background and Aims:** The atherosclerotic plaque microenvironment is highly complex, and selective agents that modulate plaque stability or other plaque phenotypes are not yet available. We sought to investigate the human atherosclerotic cellular environment using scRNA-seq to uncover potential therapeutic approaches. We aimed to make our workflow user-friendly, reproducible, and applicable to other disease-specific scRNA-seq datasets.

**Methods:** Here we incorporate automated cell labeling, pseudotemporal ordering, ligand-receptor evaluation, and drug-gene interaction analysis into an enhanced and reproducible scRNA-seq analysis workflow. Notably, we also developed an R Shiny based interactive web application to enable further exploration and analysis of the scRNA dataset.

**Results:** We applied this analysis workflow to a human coronary artery scRNA dataset and revealed distinct derivations of chondrocyte-like and fibroblast-like cells from smooth muscle cells (SMCs), and show the key changes in gene expression along their de-differentiation path. We highlighted several key ligand-receptor interactions within the atherosclerotic environment through functional expression profiling and revealed several attractive avenues for future pharmacological repurposing in precision medicine. Further, our interactive web application, *PlaqView* (www.plaqview.com), allows other researchers to easily explore this dataset and benchmark applicable scRNA-seq analysis tools without prior coding knowledge.

**Conclusions:** These results suggest novel effects of chemotherapeutics on the atherosclerotic cellular environment and provide future avenues of studies in precision medicine. This publicly available workflow will also allow for more systematic and user-friendly analysis of scRNA datasets in other disease and developmental systems. *PlaqView* allows for rapid visualization and analysis of atherosclerosis scRNA-seq datasets without the need of prior coding experience. Future releases of *PlaqView* will feature additional larger scRNA-seq and scATAC-seq atherosclerosis-related datasets, thus providing a critical resource for the field by promoting data harmonization and biological interpretation.

## Background

Atherosclerosis is a complex process involving chronic inflammation and hardening of the vessel wall and represents one of the major causes of coronary artery disease (CAD), peripheral artery disease, and stroke [1]. Rupture of an unstable atherosclerotic lesion can lead to the formation of a thrombus, causing complete or partial occlusion of a coronary artery [2]. The contribution of smooth muscle cells (SMCs) to both lesion stability and progression has recently been established by numerous groups. However, the exact mechanisms by which SMCs modulate the atherosclerotic microenvironment and whether pharmacological agents can be used to selectively counter SMC-related deleterious mechanisms are still under investigation [3–5]

Recent advances in single-cell RNA-sequencing (scRNA-seq) have enabled ultra-fine gene expression profiling of many diseases at the cellular level, including atherosclerotic coronary artery disease [5]. As sequencing costs continue to decline, there has also been a consistent growth in scRNA datasets, data analysis tools and applications [6]. Currently, a major challenge with scRNA-seq analysis is the inherent bias introduced during manual cell labeling, in which cells are grouped by clusters and their identities called collectively based on their overall differential gene expression profiles [7]. Another draw-back inherent to commonly used scRNA-seq protocols is their destructive nature to the cells, making time-series analyses of the same cells impossible. Instead, these studies must rely on time-points from separate libraries to monitor processes such as clonal expansion and cell differentiation [8,9].

Recently, new approaches have been developed to compensate for both of these shortcomings, namely automatic cell labeling, pseudotemporal analysis, and trajectory inference. Tools such as ‘SingleR’, ‘scCATCH’, and ‘Garnett’ have been used to assign unbiased identities to individual cells using reference-based and machine learning algorithms, respectively [7,10,11]. Moreover, tools such as ‘Monocle3’ and ‘scVelo’ align and project cells onto a pseudotemporal space where each cell becomes a snapshot within the single-cell time continuum [12,13]. In essence, the single scRNA-seq dataset is transformed into a time series [12–14]. Although the pseudotemporal scale does not reflect the actual time scale, it is a reliable approximation to characterize cell fate and differentiation events, e.g., during organogenesis, disease states, or in response to SARS-CoV-2 infections [13,15]

In this study, we present the application of an enhanced, scalable, and user-friendly scRNA-seq analysis workflow on an existing human coronary artery scRNA-seq dataset. We performed unbiased automatic cell identification at the single-cell level, pseudotemporal analysis, ligand-receptor expression profiling, and drug repurposing analysis. Our results demonstrate potential new mechanisms by which SMCs contribute to the atherosclerotic phenotype and signaling within the lesion microenvironment. More importantly, we revealed attractive candidate avenues for future pharmacological interventional studies. We also developed an interactive web application to allow other users to explore this dataset. This reproducible analysis pipeline and application can also be easily modified to incorporate different tissue data sources and single-cell modalities such as scATAC-seq [16] or CITE-seq [17], and could serve as a template to analyze and visualize single-cell datasets in other disease models.

## Results and Discussion

### Unbiased automatic cell labeling reveals abundant cells with chondrocyte and fibroblast characteristics

Recently, automatic cell identification tools have been introduced to compensate for the shortcomings of manual, cluster-based cell labeling [7]. For example, ‘SingleR’ and ‘Garnett’ use reference-based and machine learning algorithms, respectively, to call individual cell identities [7,10,18]. Using ‘SingleR,’ which uses known purified cell expression data as references, we found that endothelial cells (ECs) make up the highest proportion of cells in this dataset (16.21%, Figure 1A-B), followed by smooth muscle cells (SMCs, 13.8%) and stem cells (SCs, 14.06%), where the latter could be so-called “atherosclerotic stem cells” or normal stromal stem cells but cannot be distinguished until specific expression profile references are developed in the future [8]. Consistent with recent scRNA-seq studies in atherosclerotic models, we identified abundant fibroblast (FB) and chondrocyte-like (CH) cells, as well as cells with an osteoblast-like (OS) expression profile (Figure 1B) [4]. In the UMAP clusters reflecting single-cell identities, there was a substantial presence of SMC and FB cells in the OS and SC cluster. Such heterogeneity in cell clusters would have been overlooked in manual cluster-based cell labeling. We also applied another reference-based cell calling tool ‘scCATCH,’ and found that it underperforms relative to ‘SingleR’ and fails to provide consistent cell type assignment when provided with similar tissue priors (Supplementary Table 1).

**Figure 1.**
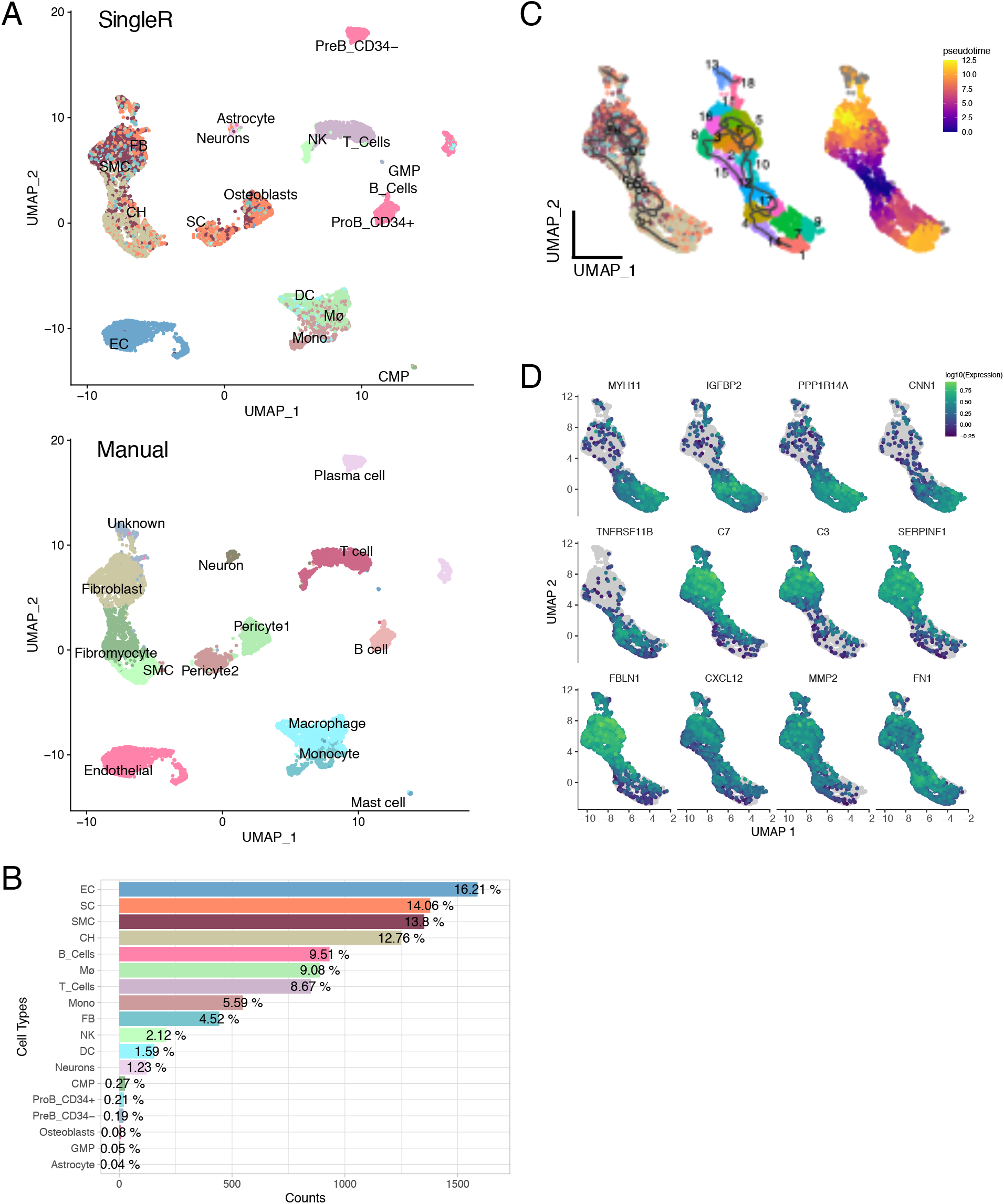
Unbiased automatic cell labeling of human coronary scRNA data using ‘SingleR’ reveals abundant cells with chondrocyte (CH) and fibroblast (FB) gene expression patterns, and pseudotemporal and ultra-fine clustering reveals derivation of CH and FB-like cells from SMCs. **(A)** UMAP clustering of 9798 cells derived from human coronary artery explants with labels based on singleR annotation (Top) and cluster-based manual annotation based on Wirka et al., 2019 (Bottom). **(B)** population breakdown by percentage. SMC: smooth muscle cells, EC: endothelial cells, CH: chondrocytes, FB: fibroblasts, Mø: macrophages, SC: stem cells, CMP: common myeloid progenitor cells, GMP: granulocyte-monocyte progenitor cells. **(C) Left**, RNA trajectory (line) shows path two direct paths from the SMC starting nodes (grey circles). **Middle**, ultra-fine clustering shows the logical transition stages from SMC to CH and FB. **Right**, pseudotemporal analysis confirms that the cell and clusters existing along a logical single-cell continuum. **(D)** selected genes that were shown to vary over pseudotime by Moran’s I test were visualized.

### Pseudotemporal ordering identifies distinct chondrocyte and fibroblast-like cell differentiation states from smooth muscle cells

To evaluate putative cell fate decisions or differentiation events (e.g., SMC phenotypic transition states), we performed pseudotemporal analysis and ultra-fine clustering using ‘Monocle3’, a method previously applied to normal and diseased states, e.g., embryo organogenesis and response to coronavirus infection, respectively [13,15]. We compared over 60 other trajectory inference (TI) methods including Slingshot [19], PAGA [20], and SCORPIUS using the ‘Dynverse’ package [21]. However, we found that most TI algorithms were unsuitable for complex tissue environments such as the atherosclerotic plaque due to their inability to distinguish disconnected topologies (Supplementary Table 2). In particular, we found evidence of SMCs giving rise to both chondrocyte (CH) and fibroblast (FB)-like cells (Figure 1C). This corroborates earlier findings showing that SMCs may transition or de-differentiate into ‘fibromyocytes’—SMCs that have undergone a phenotypic modulation to an extracellular matrix producing cell type within the atherosclerotic lesions [5,8]. Genes associated with healthy SMC phenotypes, such as *MYH11* (a canonical marker of SMC), *IGFBP2* (associated with decreased visceral fat), and *PPP1R14A* (which enhances smooth muscle contraction), are decreased by approximately 50-75% along the SMC trajectory as these cells become more FB-like (Table 1, Figure 1D, p < 0.1E-297) [3,22]. Similar results were found by another group using mouse lineage-traced models where MYH11 expression was decreased in SMC-derived modulated “intermediate cell states” [4].

**Table 1.** Selected Moran’s I statistics for genes listed in Figure 1D. Moran’s I statistics was used to identify spatial correlations within the single-cell trajectory. +1 indicates that nearby cells are perfectly similar, 0 indicates no similarity or pattern, and -1 indicates total dissimilarity [13].

|  | p_value | morans_test_statistic | morans_I |
| --- | --- | --- | --- |
| MYH11 | <1E-297 | 169.0285919 | 0.66505236 |
| IGFBP2 | <1E-297 | 129.2299117 | 0.50844307 |
| PPP1R14A | <1E-297 | 169.4172149 | 0.66663148 |
| CNN1 | <1E-297 | 170.9288702 | 0.67251583 |
| TNFRSF11B | <1E-297 | 97.66515083 | 0.38400709 |
| C7 | <1E-297 | 156.2248703 | 0.61472417 |
| C3 | <1E-297 | 103.4495 | 0.7252684 |
| SERPINF1 | <1E-297 | 157.9946249 | 0.62168635 |
| FBLN1 | <1E-297 | 154.3399713 | 0.60730135 |
| CXCL12 | <1E-298 | 48.92096 | 0.3428102 |
| MMP2 | <1E-299 | 62.57231 | 0.4385507 |
| FN1 | <1E-297 | 125.4354275 | 0.49354242 |

More importantly, specific inflammatory markers and proteins associated with thrombotic events during CAD, including complement proteins *C7* and *C3, FBLN1*, and *CXCL12* are increased along the same trajectory (Figure 1D) [2,23]. Recent evidence suggests that CAD-associated *CXCL12* secreted from endothelial cells may promote atherosclerosis[24]. Our results point to a potentially new source of *CXCL12* that could be targeted to inhibit SMC-to-FB dedifferentiation. Together, and in corroboration of recent studies, our pseudotemporal analysis demonstrates that SMCs could be a source of both FB and CH-like cells, associated with intermediate and advanced atherosclerotic phenotypes, respectively [4,5]. This is further supported by a recent study, in which blocking of SMC-derived intermediate cells coincides with less severe atherosclerotic lesions [4]. Precisely how these cells might influence the overall stability of atherosclerotic lesions and clinical outcomes requires additional longitudinal studies using genetic models and deep phenotyping of human tissues [3,4,8].

### Comprehensive ligand-receptor analysis shows complex intercellular communications in the human coronary micro-environment and reveals potential drug targets

To examine the potential cross-talk between different cell types using scRNA-seq data, we compared the ligand and receptor expression profiles of each cell type with experimentally-validated interactions using ‘scTalk’ [25]. We found that there is an intricate network of signaling pathways connecting different cell types; some cell types, such as OS, have stronger and more frequent outgoing signals, whereas other cell types such as Macrophages (Mø) have fewer and weaker incoming and outgoing signals (Figure 2A). SMCs, OSs, and neurons also exhibit a high degree of autocrine signaling profiles (Figure 2A). Specifically, SMCs are shown to have the highest number of outgoing signals and are among those with the least number of incoming signal weights (Figure 2B). This suggests that SMCs play an important role in regulating the coronary microenvironment by transducing signals to neighboring cells in the lesion.

**Figure 2.**
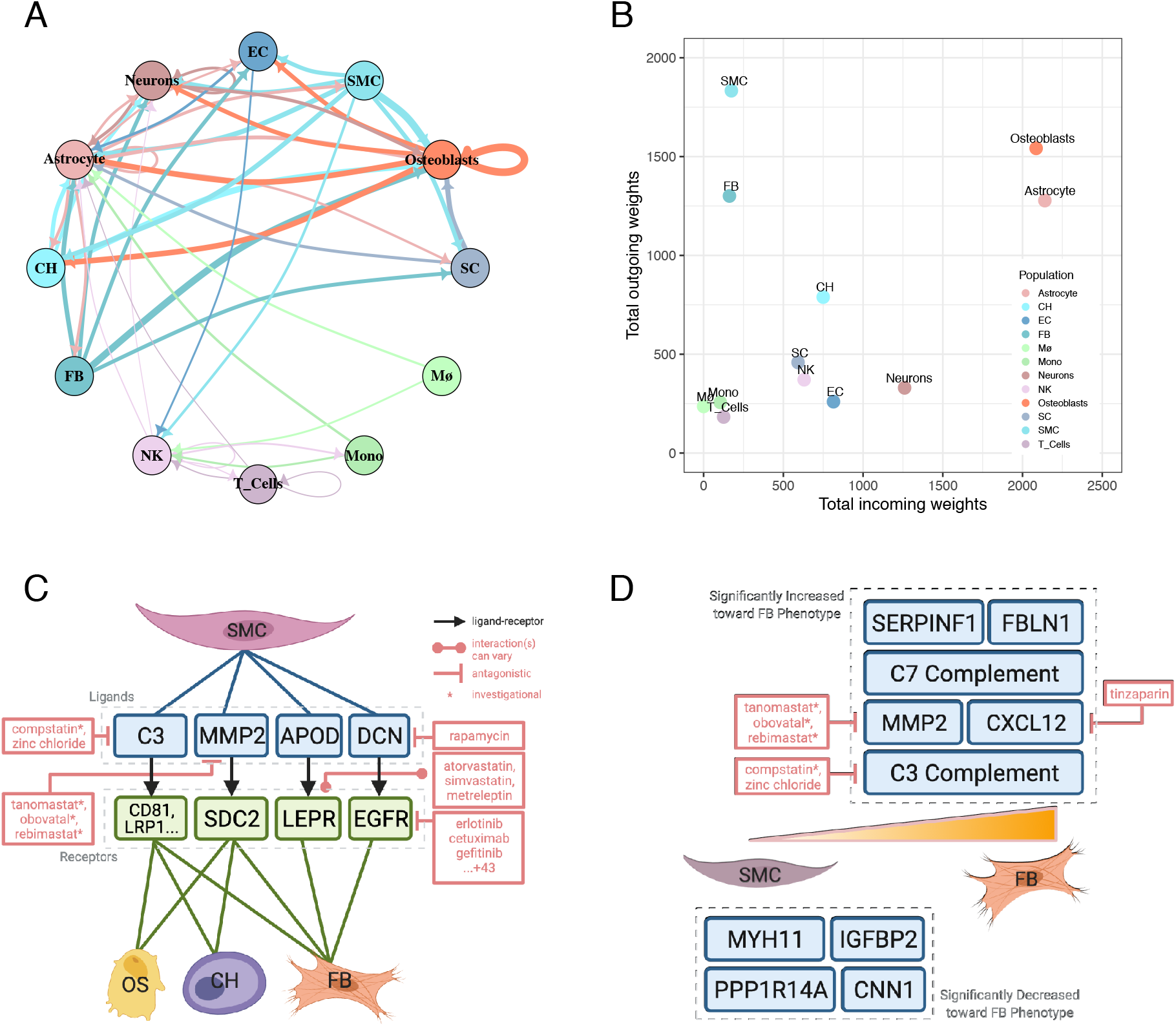
Comprehensive ligand-receptor analysis coupled to drug-gene interaction analysis reveals potential drug therapies to interrupt SMC to CH and FB communication and SMC trajectory. **(A)** circle plot representation of the inferred intercellular communications within the coronary artery environment. **(B)** incoming and outgoing signal summed weights by cell type. **(C)** ‘scTalk’ shows that SMCs interacts with FB through three pairs of experimentally verified interactions. Of these three, complement and *DCN*-*EGFR* signaling is the most druggable as revealed by DGIdb 3.0. **(D)** druggable genome analysis revealed agents that can target genes such as complement signaling (C3), C-X-C Motif Chemokine Ligand 12 (CXCL12), and matrix metalloprotease (MMP2), which increases along the SMC-to-FB phenotype trajectory.

Specifically, ten signaling pathways were identified between SMC, FB, CH, and OS (Supplemental Figure 1). These pathways involve C3 complement, fibulin-1 *(FBLN1)*, apolipoprotein D *(APOD)*, decorin *(DCN)*, and matrix metalloprotease 2 (*MMP2*, Figure 2C, Supplemental Figure 1). We searched for potentially druggable targets to interrupt SMC-FB communication by performing an integrative analysis of the identified ligand-receptor interactions and the druggable genome using several drug-gene interaction databases, including DGIdb 3.0 and Pharos [26]. We found that SMCs signals to FBs through the complement protein *C3* and syndecan-2 *(SDC2)* via *MMP2*, and these two pathways can be disrupted by drugs such as compstatin and tanomastat, respectively (Figure 2C). Multiple studies have linked *C3* and the complement system to atherosclerotic lesion maturation in mouse models, and a recent case study showed that the C3 targeted inhibitor, compstatin (AMY-101), may prevent cardiovascular complications in patients with severe COVID-19 pneumonia [27–30]. Our results provide a potential mechanistic explanation by which SMCs can modulate the inflammatory environment and plaque formation. Further, a recent study demonstrated that *microRNA-9* repression of *SDC2* impedes atherosclerosis formation [31], while MMP2 alteration also contributes to atherosclerosis in mouse models [32]. Here, we show that SMCs signal to FBs within the atherosclerotic environment via *SDC2*-*MMP2*, and reveal additional upstream candidate drug therapies that may influence atherosclerosis progression.

Interestingly, anti-*EGFR* (epidermal growth factor receptor)-based cancer treatments such as erlotinib, cetuximab, and gefitinib were identified as potential key mediators of signaling pathways between SMCs and FBs via decorin (*DCN*) and *EGFR* (Figure 2C). It has been shown that *DCN* overexpression increases SMC aggregation and SMC-induced calcification within atherosclerotic plaque [33]. Although the overlap between CAD and cancer etiology has been previously noted, the long-term efficacy and cardiovascular impact of chemotherapy drugs, such as erlotinib, requires further translational studies to investigate their potential use in cancer patients to treat CAD [34–36].

Furthermore, integrative analysis of gene expression variation along the SMC-to-FB RNA trajectory revealed similar results; as shown earlier in Figure 1D, the expression of complement genes such as *C3* and *C7*, and chemokine *CXCL12* are increased as SMCs become more FB-like. Although CXCL12 derived from endothelial cells have been recognized to promote CAD in mouse models [24], here we provide a potentially new source of CXCL12 using human data and found several pharmacological agents such as tinzaparin, an FDA-approved anticoagulant, to investigate in future interventional studies. Together, this combination of independent analyses of a scRNA-seq dataset reaffirms the druggable potential of these target genes.

### *PlaqView* is a user-friendly web application to explore atherosclerosis-related datasets

To enable other researchers to explore the transcriptomic landscape of the atherosclerotic environment, we developed a web interface called *PlaqView* (www.plaqview.com, Figure 3A). This interactive, R Shiny-based tool allows for multiple gene queries and comparisons of gene expression, cell-labeling methods, RNA-trajectory tools, integrative drug-gene analysis, and outputs high quality graphs and detailed tables. To our knowledge, there are no publicly available tools to visualize atherosclerosis-related single-cell datasets without prior coding knowledge. Further, *PlaqView* is under active development and will be releasing new datasets coincidently with future atherosclerosis-related publications. As the database in *PlaqView* expands along with the growing number of single-cell datasets, we anticipate that it will become an essential tool for the atherosclerosis research field.

**Figure 3.**
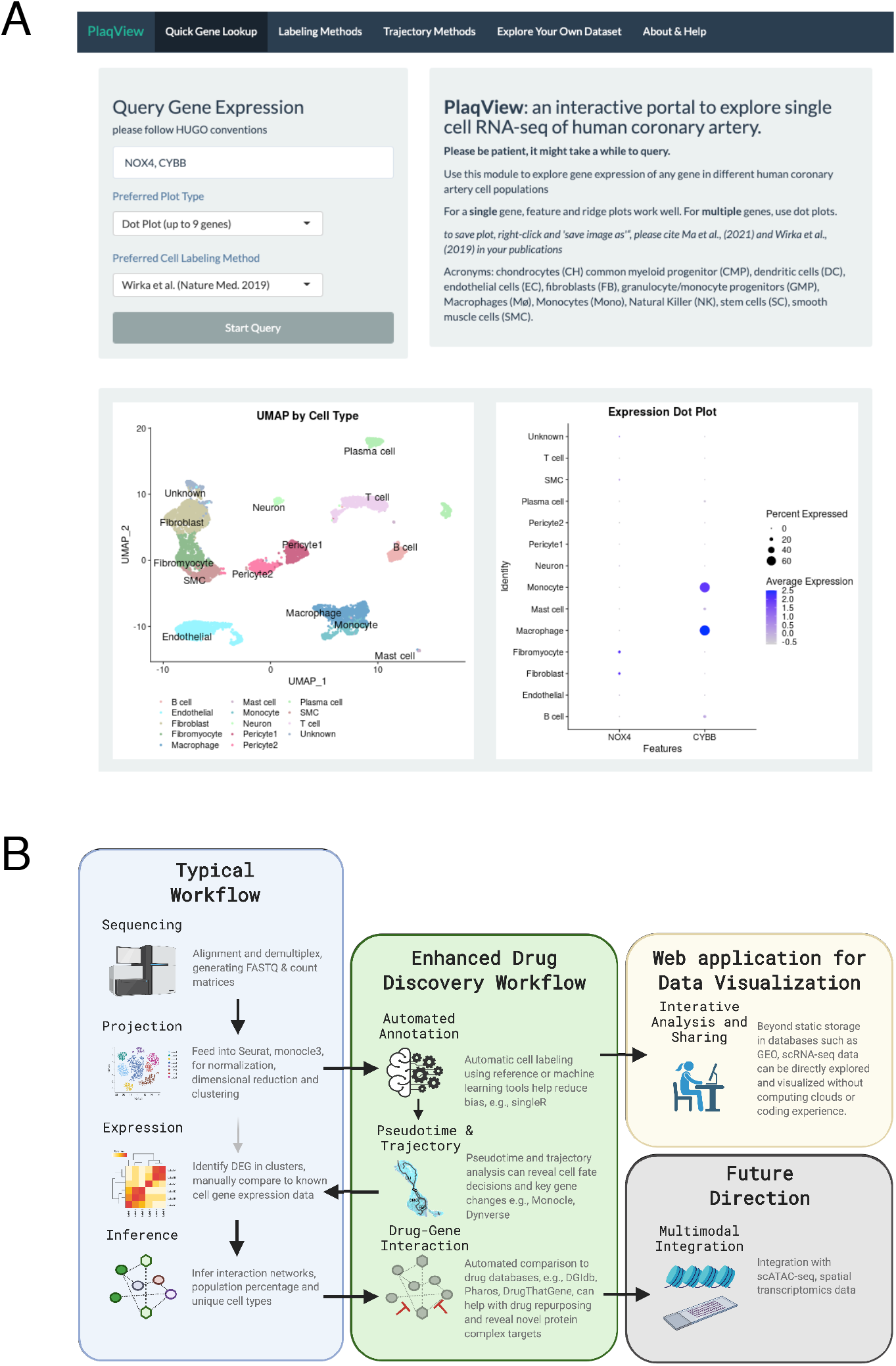
**(A)** screenshot of the PlaqView web application. **(B)** a roadmap of enhanced scRNA-seq analysis process. Instead of cluster-based grouping, our pipeline uses automatic cell labeling, coupled with pseudotime trajectory, cellular network interaction and drug targeting, and provides a reproducible process for scRNA datasets. From this roadmap, it is easy to add additional analysis tools and modify workflow as more tools and datasets become available.

### Limitations

Despite the advances presented in this workflow, we are still working to improve several limitations. For instance, the default reference in ‘SingleR’ cannot identify more recently discovered cellular phenotypes such as fibromyocytes [5,7,37], which may require a combination of manual and automated labeling methods for accurate identification. Factors such as the intrinsic heterogeneity of the tissue sample, disease stage, and tissue processing artifacts are difficult to isolate computationally and could influence automated labeling methods. Nonetheless, interactive viewers and tools such as *PlaqView* which incorporate multiple methods and visualizations in one location, could help users separate out the technical and biological variation from various single-cell datasets. Additionally, the modular nature of *PlaqView* will also allow for future improvement of labeling methods as more precise reference datasets are made available. Lastly, the true efficacy of proposed drugs cannot be verified without extensive pre-clinical testing and clinical trials. Nonetheless, these findings may catalyze future investigative efforts to develop more targeted therapies.

## Conclusions

Our findings show that an enhanced, reproducible pipeline for scRNA-seq analysis has the potential to improve upon current standard scRNA-seq bioinformatics protocols. For instance, we provide new insights into intricate vascular cell differentiation and communication pathways while providing actionable and testable targets for future experimental studies (Figure 3B). In our combined analysis, we found that SMCs give rise to substantial proportions of CH and FB, with the latter associated with worse prognostic markers [2,4,22,23]. SMCs signal to FBs via a series of pathways involving C3 complement and MMP2, whose expression coincidentally increases along the SMC-to-FB trajectory. We revealed possible therapeutic avenues that may disrupt these cell communication pathways and alter the atherosclerotic pathology. Furthermore, several FDA-approved drugs (e.g., erlotinib, cetuximab, and gefitinib) were shown as potential effectors of SMC signaling to FB, and may be used to treat CAD in cancer patients to simplify or augment drug regiments [34]. This is consistent with recent reports showing beneficial effects of the acute promyelocytic drug all-trans-retinoic acid (ATRA) in atherosclerosis mouse models [4]. Further investment in scRNA-seq may also help resolve the balance of anti-tumor efficacy and atheroprotection for immune checkpoint inhibitors as well as immunomodulators at the interface of cardio-immuno-oncology [38].

Although the utilization of this workflow can compensate for many of the shortcomings of current scRNA-seq analyses, we are still unable to perform cell-lineage tracing that reflects actual timescales without additional gene engineering experiments *in vivo* [39]. However, leveraging mitochondrial DNA variants in snATAC-seq data has enabled lineage tracing analysis in human cells [40,41]. Likewise, these analyses can ultimately be extended to integrate spatial omics and other multi-modal data [42]. In the future as spatial transcriptomics, scATAC-seq, and/or CITE-seq data become more widely available, this workflow can be modified to discover signaling pathways or differentiation events at specific tissue locations and timepoints, allowing for more disease-relevant drug-gene interaction analyses (Figure 3B). Nonetheless, this pipeline can be applied immediately to datasets from other tissues/diseases to generate informative directions for follow-up studies, and is more user-friendly and reproducible compared to standard scRNA analyses. Lastly, building web applications that democratize the access and analysis of single-cell data will promote collaboration and innovation across disciplines [43]. As *PlaqView* incorporates additional relevant single-cell datasets, we anticipate that this application will become an indispensable resource for the community.

## Methods

### Data retrieval and pre-processing

Human coronary artery scRNA data read count matrix was retrieved from the Gene Expression Omnibus (GEO) using #GSE131780 and loaded into R 4.0, and was preprocessed using standard parameters of the R packages ‘Seurat’ v.3, and ‘Monocle3’ as required [5,44–46]. Uniform manifold approximation projections (UMAP) clusters from ‘Seurat’ were imported into ‘Monocle3’ before pseudotemporal analysis.

### Automatic cell Identification and pseudotemporal ordering

scRNA read matrices were read into SingleR as previously described for cell labeling [7]. SingleR compares each cell’s gene expression profile with known human primary cell atlas data and gives the most likely cell identity independently. SingleR first corrects for batch effects, then calculates the expression correlation scores for each test cell to each cell type in the reference, and the cell identity is called based on reference cell type exhibiting the highest correlation.

Then, pseudotemporal analyses were performed as previously described in the analysis of embryo organogenesis [13,45]. Briefly, the UMAP clusters were passed into Monocle3 and then the ‘learn_graph()’ and ‘order_cells()’ functions. The SMCs and related clusters were then subsetted for detailed subclustering and analysis. For each cluster, Moran’s I statistics were calculated, which identify genes that are differentially expressed along their trajectories. Additionally, we explored 60+ other trajectory inference methods using the ‘Dynverse’ package, where we simulated mock dataset of 5000 cells with known disconnected graph topologies [21]. Detailed codes to reproduce the figures in this publication can be found at the Miller Lab Github (see availability of data and materials).

### Ligand-receptor cell communication analysis

We analyzed candidate ligand-receptor interactions to infer cell communication using the R package ‘scTalk’, as previously described in the analysis of glial cells [25]. This method is based on permutation testing of random networks, where ligand-receptor interactions are derived from experimentally derived interactions from the STRING database. We exported statistically significant differentially expressed genes from ‘Seurat’ using the ‘FindMarkers()’ function and imported the preprocessed data into ‘scTalk.’ Then, overall edges of the cellular communication network were calculated using the ‘GenerateNetworkPaths()’ function, which reflects the overall ligand-receptor interaction strength between each cell type. Then, the cell types of interest were specified and treeplots were generated using the ‘NetworkTreePlot()’ function.

### Gene-drug interaction analysis

The above identified ligand and receptor interaction pairs were fed into the Drug-Gene Interaction database (DGIdb 3.0) to reveal candidate drug-gene interactions [26]. Ligands and receptors that were deemed significant from ‘scTalk’ were evaluated using the ‘queryDGIdb()’ function of the ‘rDGIdb’ R package [26]. Additionally, we queried CTD2, OMIM, ClinVar, Pharos, GnomAD, and the ExAC databases using the docker-based tool ‘DrugThatGene’ [47]. We included all top FDA-approved drugs produced with verified inhibitory or antagonistic activities, as well as drugs that may influence or influenced by changes in the receptor. Figures 2C, 2D, and 3B were modified using BioRender for clarity.

### Development of *PlaqView*

*PlaqView* is written in R and Shiny, and is hosted on the web at www.plaqview.com using dedicated shinyapps.io servers, and can be run locally through RStudio. The raw data was first processed as previously described, but packaged into .rds objects. These objects were then written into a Shiny script to allow for interactive display. This application is open-sourced and its code and data is available at github.com/MillerLab-CPHG/PlaqView. Further, *PlaqView* is actively recruiting available datasets on the github page.

## Declarations

No conflicts of interest to disclose.

## Ethics approval and consent to participate

Not applicable

## Consent for publication

Not applicable

## Availability of data and materials

All data are publicly available on GEO, accession number GSE131778, as previously described [5]. All codes and analysis pipelines can be viewed at github.com/MillerLab-CPHG/Ma_2020 and repurposed for other scRNA datasets.

## Authors’ contributions

WFM designed and performed the statistical analysis. CJH, AWT, DW and YS refined the methodology and edited the manuscript. JVM helped with scripting. CLM conceived and refined the project and edited the manuscript.

## Acknowledgements

Not applicable.

## Abbreviations

C7: complement component C7
CAD: coronary artery disease
CH: chondrocytes
CMP: common myeloid progenitor cells
DCN: decorin
DGIdb: drug-gene interaction database
EC: endothelial cells
EGFR: epidermal growth factor receptor
FB: fibroblasts
FBLN1: fibulin 1
GMP: granulocyte-monocyte progenitor cells.
Mø: macrophages
MYH11: myosin heavy chain 11
SC: stem cells
sc/snATAC-seq: single cell/single nucleus assay for transposase-accessible chromatin sequencing
sc/snRNA-seq: single cell/single nucleus RNA sequencing
TI: trajectory inference
SMC: smooth muscle cells
UMAP: uniform manifold approximation and projection

