## Supplementary figures and images for "Single-cell RNA-seq analysis of human coronary arteries using an enhanced workflow reveals SMC transitions and candidate drug targets"

### Supplemental Figure 1

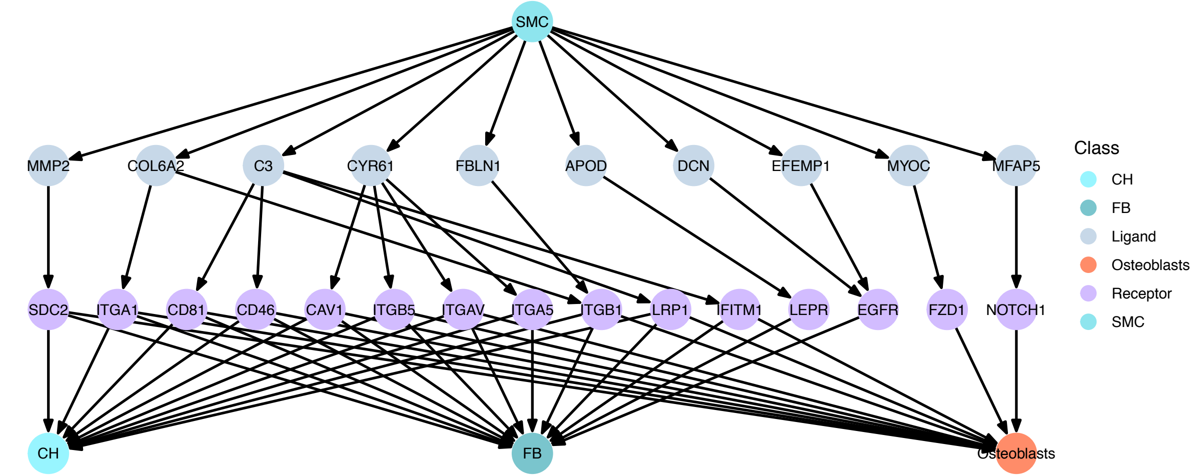
